# Patching up the nucleus: a novel role for PMLII in nuclear envelope rupture repair

**DOI:** 10.1101/2025.01.24.634656

**Authors:** Anne F.J. Janssen, Oliver Knowles, Sébastien Britton, Janet E. Deane, Evan Spruijt, Delphine Larrieu

## Abstract

The nuclear envelope (NE) is important for cellular health as it protects and organizes the genome. NE dynamics is important for various cellular processes including cell growth, migration and removal of defective NE components. In extreme cases, the NE can rupture leading to exchange of material between the nuclear interior and the cytoplasm. Rapid repair of the NE is initiated to minimize the effect on the genome. While our understanding of the machinery involved in this repair process is increasing, a lot is still unknown about this process including events leading up to NE rupture. Interestingly, biomolecular condensates have recently been found to play important roles in membrane repair and remodelling in cells. Here, we found that promyelocytic leukemia protein isoform II (PMLII), a protein involved in nuclear PML body formation, forms condensates at the NE. These condensates specifically form at sites where the lamina is disrupted. We show that NE rupture often occurs at these sites and that PMLII stays present until rupture repair is initiated suggesting a role in stabilization of the site for effective repair.

## Introduction

The nuclear envelope (NE) plays important roles in organizing and protecting the genetic information of the cell, transducing signals and regulating trafficking between the cytoplasm and the nucleus. The NE is composed of an outer (ONM) and inner nuclear membrane (INM) which are connected at sites of the nuclear pore complex (NPC). The lamina, a filamentous network of A type (lamins A and C) and B type (lamin B1 and B2) intermediate filament proteins, underlies the INM at the nucleoplasmic side. The lamina interacts with heterochromatin and proteins embedded in the INM and thereby plays important roles in regulating nuclear size, shape and stiffness^1,2^. Together, these structures are essential for proper cellular function as reflected by a subset of conditions, together named envelopathies, caused by mutations in NE proteins. These diseases include muscular dystrophies^3^, dilated cardiomyopathy^4^ and premature aging syndromes^5–7^. Moreover, nuclear envelope dysfunction has been correlated with cancer^8^, aggregation diseases and general aging^9^.

The NE can undergo remodelling during interphase which is important for various processes including cell growth, differentiation, NPC insertion, removal of defective NE components and nuclear egress of larger particles^10^. In extreme cases, NE rupture can occur, resulting in loss of nuclear compartmentalization^11,12^. NE ruptures are typically observed in cells with defective lamina integrity caused for example by loss or mutations of NE proteins, or in cells experiencing mechanical stress, e.g. during confined migration. Recent efforts have been focussed on increasing our understanding of how NE ruptures are repaired and what the functional consequences of NE rupture are^11–18^. However, much is still unknown about the regulation of NE and lamina remodelling events including those that precede a rupture. How do lamin holes form? Are there proteins marking sites of disrupted lamina? What initiates membrane remodelling at bleb sites?

Intriguingly, biomolecular condensates have recently been shown to be involved in membrane remodelling events, intracellular transport and membrane repair^19–22^. Biomolecular condensates form through phase separation of biomolecules with weak multivalent interactions. They can concentrate biomolecules and are now recognized to play a role as reaction hubs or storage compartments in cellular organizations^23,24^. The nucleus contains various biomolecular condensates including the nucleolus, PML bodies and nuclear speckles.

Here we describe that promyelocytic leukemia protein isoform II (PMLII), a PML body constituent, can form condensates at the NE. These PML patches are found at locations where the lamina is weakened and are therefore often sites of NE rupture events. We describe that the unique PMLII C-terminus, predicted to be mainly disordered but containing two amphipathic α-helices, is important for this specific localization. Finally, we find that the PML patch starts disassembling after initiation of NE repair suggesting a role in stabilizing the rupture site allowing membrane repair to occur.

## Results

### PMLII localises to holes in the lamina

NE ruptures often occur at locations where the lamina is weakened. These gaps in the lamina, or lamin holes, are generally found to be lacking many NE proteins such as LEM domain proteins, that are embedded in the INM and nuclear pore complex proteins (NUPs). While looking for proteins present in these areas, we found that endogenous PML proteins, the key organizer of the PML nuclear body, can localize to lamin holes in multiple cell lines (Fig 1a, arrows). This localization was already described by others in overexpression conditions and found to be a specific property of PML isoform II^25,26^. The PML gene contains 9 exons of which the first four, containing the RBCC (RING, B-Box, Coiled Coil) motif are present in all isoforms. The RBCC motif contains multiple structurally conserved domains capable of dimerization and multimerization contributing to formation of PML bodies. Alternative splicing of the C-terminus results in 7 main isoforms: PMLI-VII, with distinct roles and binding partners^27^. PML body biogenesis is complex and incompletely understood but current models suggest a combination of oxidative stress induced cross-linking of PML monomers, oligomerization through weak non-covalent interactions in the RBCC domain, SUMO-SIM interactions and possible contributions from weak interactions between (disordered) C-termini^28^.

**Figure 1:**
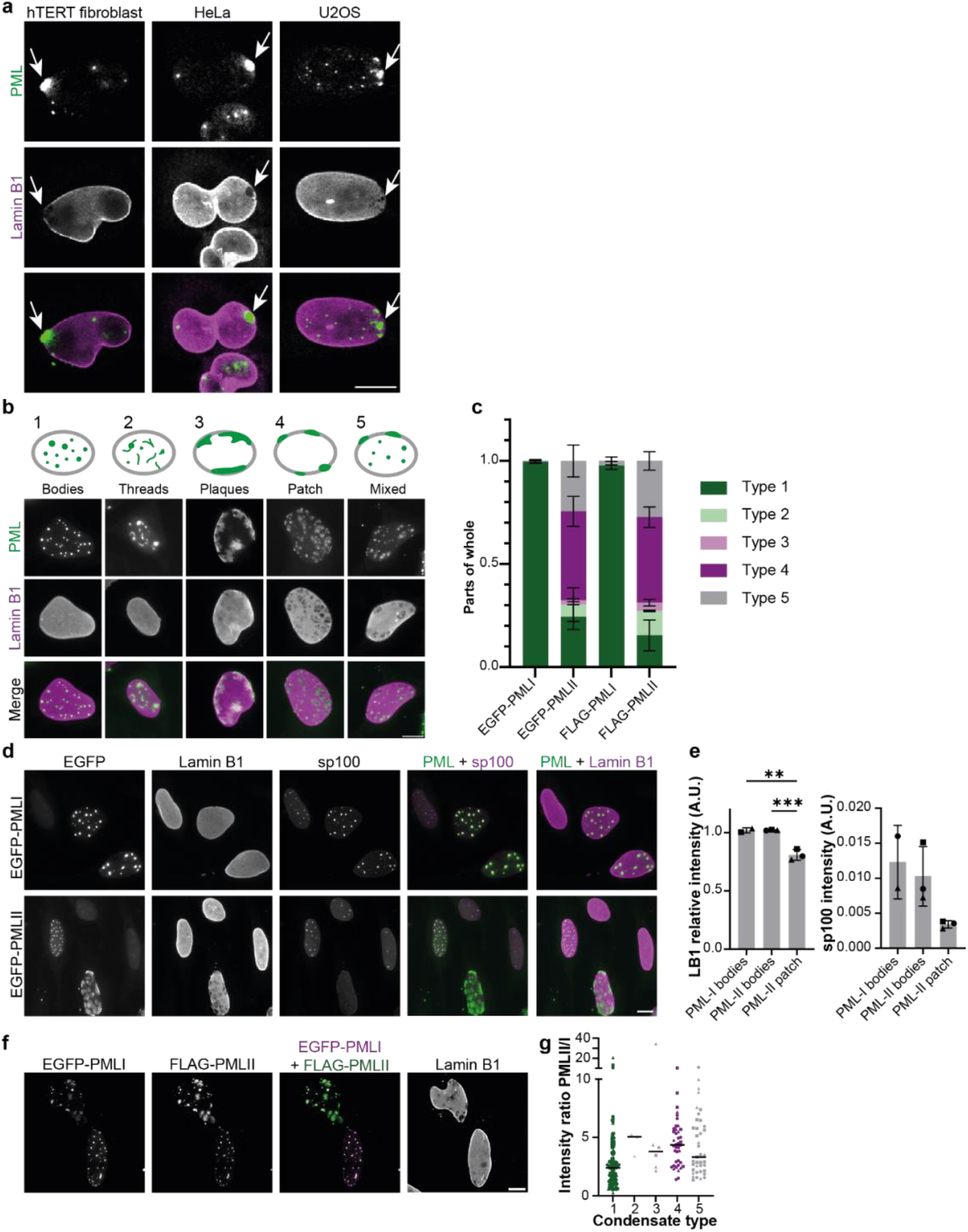
PMLII localizes to holes in the lamina. **a)** Representative lamin B1 and PML confocal immunofluorescence images from human cell lines. The arrows indicate areas of PML accumulation at NE areas devoid of lamina. Scale bar: 10 μm. **b)** Different condensate types observed upon overexpression of EGFP-PMLII in HeLa cells including widefield immunofluorescent images of lamin B1. Scale bar: 10 μm. **c)** Quantification of condensate types observed for PMLI or PMLII overexpression in HeLa cells. N=3, for EGFP-PMLI and EGFP-PMLII and N=2 for FLAG-PMLI and FLAG-PMLII. Mean +/-sd is indicated. **d)** Representative, lamin B1 and sp100 widefield immunofluorescence images of HeLa cells overexpressing EGFP-PMLI or EGFP-PMLII. Scale bar: 10 μm. **e)** Quantification of relative lamin B1 intensity and sp100 intensity at puncta or patches in HeLa cells overexpressing EGFP-PMLI (N=2 (n=18, 32 cells)) and EGFP-PMLII (puncta; N=3 (n=21,14,13 cells), patches; N=3(n= 11,17,17 cells)). Mean +/-sd are indicated, independent experiments are indicated by the different symbols. P-value was calculated using a one-way ANOVA test followed by Tukey’s post hoc test (**P<0.01, ***P< 0.001). **f)** Representative, lamin B1 confocal immunofluorescence images of HeLa cells co-overexpressing EGFP-PMLI and FLAG-PMLII. Scale bar: 10 μm. **g)** Quantification of PMLI/PMLII intensities per condensate type. n=112, 3,6,40, 39 for types 1-5 respectively. Data are presented from three independent experiments indicated by the triangels, squares and cirkels. Median is indicated.

### PMLII forms different condensate types in cells

We first investigated the difference in condensate formation between the two main isoforms of PML, PMLI and PMLII^29^. We found that while EGFP-PMLI only forms punctate condensates inside the nucleus, the classical PML bodies reported before, EGFP-PMLII can form a variety of condensates. We distinguished four condensate types: spherical PML bodies inside the nucleus (type 1), amorphous/thread like bodies (type 2) inside the nucleus, irregularly shaped NE-localized agglomerates hereafter referred to as plaques (type 3), and round NE-localized patches at holes in the lamina (type 4) (Figure 1b). Manual classification showed that PMLII mainly forms patches (type 4) and bodies (type 1) (Figure 1b-c). Thread like bodies (type 2) and plaques (type 3) were only observed in 6.0% and 2.1% of cells respectively (Figure 1c). We also noticed that although plaques seem to localize to the NE, they localize to areas where the lamina is present in contrast to patches that localize to lamin holes. Cells can also display a mixture of condensate types that we classified separately (type 5). Finally, we did not observe different distributions of body types when overexpressing FLAG or EGFP tagged PMLII (Figure 1c), confirming that this is not an artifact caused by the tag itself.

We decided to focus on patch and body type condensates as these were the most frequent condensate types observed upon PMLII overexpression and were observed when staining endogenous PML. We wondered whether PMLII patches resembled PML bodies in their capacity to recruit client proteins, or whether these patches constitute a compositionally different type of PML condensate. We looked at sp100, one of the first identified client proteins of PML bodies that is recruited to PML bodies via SUMO-SIM interactions^30,31^. PML proteins and clients of PML bodies are heavily sumoylated and in addition often carry a Sumo Interaction Motif (SIM). Although SUMO-SIM interactions are not essential for PML body formation they affect PML body structure, dynamics and likely function. We found that PML bodies formed by overexpression of PMLI and II were comparable in their ability to recruit sp100 while PMLII patches seemed to recruit 3x less client protein compared to bodies (Figure 1d, e). Therefore, it seems that PMLII patch composition differs from that of PMLII bodies, suggesting these patches might perform a different function. Finally, we confirmed that PMLII patches localize to areas of low density lamina, while PML bodies (type 1) formed by either PMLI or II do not (Figure 1d, e).

When performing these overexpression experiments using the PMLI and PMLII isoform, we noticed that PMLII overexpression resulted in the formation of many PML patches, something not typically observed under endogenous conditions. We wondered whether the balance of PML isoforms could play a role in regulating PML condensate type formation. We therefore co-overexpressed PMLI and PMLII. We determined the ratio of PMLII/PMLI intensity in cells and classified PMLII condensate type according to the early identified types. We found that cells with a low PMLII/PMLI ratio (<1), were only able to form PML bodies (type 1), while cells with higher PMLII/PMLI ratios were able to form different condensate types (Figure 1f, g). These results suggest that PMLII can be recruited to PML bodies by other isoforms.

### The PMLII C-terminus and other properties essential for patch formation

The unique C-termini of PML isoforms provides them with the ability to interact with different binding partners, conferring unique functionality^27^. We wondered if the PMLII C-terminus, composed of exon 7b, is important for localization to PMLII patches at the NE. Structural prediction of this unique domain using AlphaFold2 results in a mainly unstructured protein with two predicted α-helices (Figure 2a, b). We first generated several truncations of the full PMLII protein by removing progressive regions of the C-terminal end (Figure S1b). We expressed these truncated forms in HeLa WT and PML KO cells and quantified the fraction of cells with the condensate types defined before (Figure 1b). When we first removed the C-terminal disordered part, Δ1, we increased the fraction of cells forming PMLII patches (type 4, Figure S1a, c). When additionally removing helix2 (Δ2), we see a complete switch in condensate type where most cells now form plaque condensates (type 3) at the NE colocalizing with the lamina (Figure S1a, c). These results suggested that helix2 is important for the formation of PMLII patches. However, additional removal of the disordered region between the helices, resulting in Δ3, surprisingly restored the ability of patch formation at lamin holes (Figure S1a, c). Finally, removal of helix1 (Δ4) resulted in formation of large PML bodies in the nuclear interior. These results indicated that both helices likely contribute to the ability of PMLII to form patches and that exposure of either helix by removing the neighbouring disordered region reinforces this ability (Δ1 and Δ3). To confirm that both helices are important, we additionally generated deletion mutants of the helices (Δhelix1, Δhelix2). While these helix deletions resulted in the preferential formation of different condensate types, both resulted in the loss of patch formation (Figure S1a, c).

**Figure 2:**
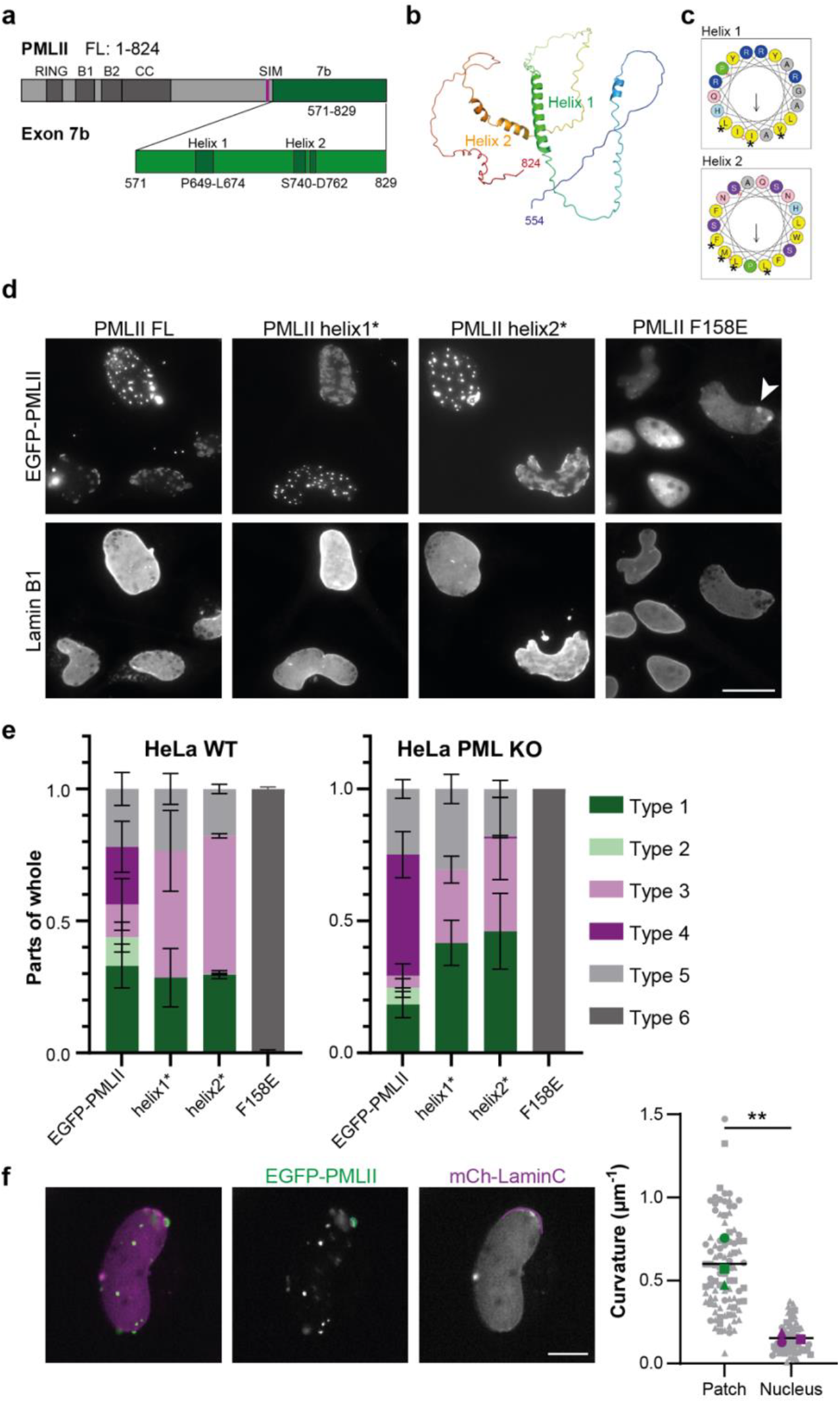
Predicted amphipathic α-helices in PMLII are important for localization to lamin holes. **a)** Schematic representation of PMLII organization with the common N-terminal RBCC domain and unique C-terminus (exon 7b). **b)** Alphafold2 prediction of PMLII C-terminus with unstructured regions and two α-helices. Coloured from blue at the N-terminal end to red at the C-terminus. **c)** α-helical wheel plot of predicted α-helices by HeliQuest server. Hydrophobic residues are shown in yellow, serine and threonine in purple, basic residues in dark blue, asparagine and glutamine in pink, alanine and glycine in grey, histidine in light blue and proline in green circles. Arrows represent direction and magnitude of the hydrophobic moment. Asterisks indicate the hydrophobic residues mutated to lose the amphipathic nature of the helices resulting in helix1* and helix2*. **d)** Representative lamin B1 immunofluorescence images of HeLa cells overexpressing EGFP-PMLII WT and mutants to lose the amphipathic nature of the helices (helix1* and helix2*) or the F158E mutant where B1 box oligomerization is impeded. Arrowhead indicates accumulation at lamin holes among mostly diffuse signal. Scale bar: 20 μm. **e)** Quantification of fractions of HeLa WT and PML KO cells overexpressing EGFP-PMLII WT and indicated mutants showing different types of condensates. Mean +/-sd indicated from N=3 independent experiments. **f)** Selected images from timelapse imaging of HeLa cells expressing EGFP-PMLII and mCherry-laminC used for curvature quantification of PMLII patches and nucleus. Scale bar: 10 μm. Example of curves that were fitted are indicated in green and magenta for patches (n=30,28,37) and nucleus (n=16,21,22) respectively. Individual datapoints are indicated in small grey circles, squares and triangles respectively and mean of N=3 independent experiments are indicated with experimental means indicated by the bigger green (patches) and magenta (nucleus) shapes. **P<0.01 using a two-tailed unpaired t-test.

As PML isoforms co-assemble through multiple interactions within their RBCC domain, we wondered whether presence of other PML isoforms may play a role in formation of different condensate types. We therefore generated a PML KO cell line (Figure S2). The results for all these truncations did not seem to be dependent on the presence of endogenous PML as results in WT and KO HeLa were similar (Figure S1c) indicating that patches and other condensate types can form by PMLII alone.

As SUMO-SIM interactions are emerging as important regulators of condensate dynamics and function^32^, we wondered whether sumoylation might affect the types of condensates formed by PMLII. We therefore mutated one of the main sumoylation sites on PMLII, generating K160R. Loss of sumoylation on Lysine-160 led to an increase of PML bodies and threads, and decreased patch formation from 43% to 3.1% of cells (Figure S1a, c). This suggest that tuning of the interactions between PML proteins or interactions with specific partners through SUMO-SIM interactions contribute to patch formation.

As both α-helices are important for patch formation, we decided to have a closer look at their properties and noticed that these were amphipathic in nature. Amphipathic helices can often interact with membranes by insertion of the hydrophobic side of the helix into the membrane^33^. To determine the importance of the amphipathic nature of the helices we mutated key residues at the hydrophobic side to polar or charged residues (Figure 2c, asterisks). Expression of these mutants, helix1* and helix2*, resulted in the complete loss of the ability to form PMLII patches, confirming that the amphipathic nature of these helices is essential for patch formation (Figure 2d,e). Insertion of α-helices into membranes often results in membrane curvature sensing or inducing properties. We indeed observed that PMLII patches often seemed to favour a curved surface, and we therefore analysed the curvature of PMLII patches. This confirmed that patches typically have a higher curvature than the curvature of the nucleus (Figure 2f).

Next, we wondered whether formation of PMLII patches was similar to body formation, where oligomerization of the B1 Box is essential for body formation^34^. Therefore, we generated the PMLIIF158E mutant carrying a point mutation in the B1 box. We found that the F158E mutant was unable to form PMLII condensates of any type leading to a diffuse localization (type 6) indicating that similar oligomerization steps are involved (Figure 2d, e). We did find that PMLII-F158E was able to show some enrichment at patch sites in WT HeLa cells (Figure 2d, arrowhead), likely caused by interactions between the PMLII C-terminus and endogenous PMLII at these patches^35^ as we did not observe enrichment in PML KO cells.

### Dynamics of PMLII patches

PMLII patches look different from PMLII bodies with a less intense PMLII signal as it seems to form a layer at the NE while PMLII bodies have bright fluorescence signal indicating a higher concentration of PMLII. We wondered whether these morphological differences also translated to different dynamics of PMLII within these condensates. Using FRAP to compare the dynamics of PMLII within patches versus bodies we found that PMLII in these different condensate types recovers equally fast and thus PMLII mobility in patches and bodies is the same (Figure 3a, b).

**Figure 3:**
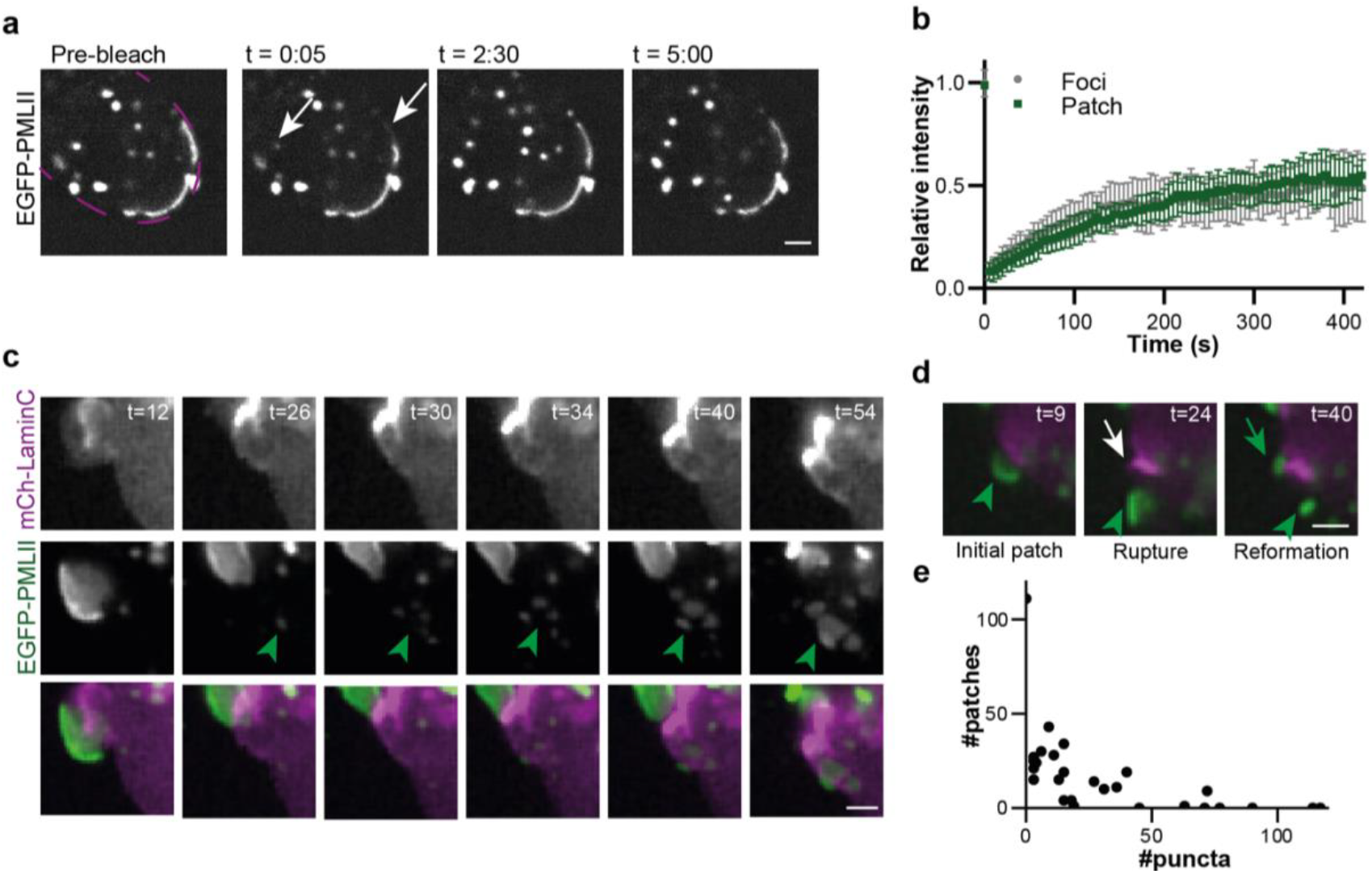
PMLII patches form slowly and show similar dynamics to PMLII puncta. **a)** Selected images from timelapse imaging from a fluorescent recovery after photobleaching (FRAP) experiment showing pre-bleach intensities and recovery of fluorescence. Time indicated is relative to time of bleaching in min:sec. Nuclear contour is indicated in magenta. Scale bar: 2 μm. **b)** FRAP recovery curves of PMLII puncta and patches. Mean +/-sd n= 7 an n=8 for puncta and patches respectively. **c)** Selected images from a time-lapse movie of HeLa cells expressing EGFP-PMLII and mCherry-laminC showing formation of PMLII patches (green arrowhead). Time is indicated in minutes. Scale bar: 2 μm. Example images from a time-lapse movie of HeLa cells expressing EGFP-PMLII and mCherry-laminC showing patch reformation. Initial patch (green arrowhead) is displaced and forms a puncta when a rupture and repair event is detected (white arrow). This is followed by reformation of a patch next to the repair site (green arrow). Time is indicated in minutes. Scale bar: 2 μm. **e)** Number of patches versus puncta in HeLa PML KO cells expressing EGFP-PMLII. n= 27 nuclei.

We then focussed on the dynamics of patch formation, growth and collapse. Although patches can form at locations all over the nucleus, we mostly observed patch formation near sites of previous rupture events (Figure 3d) indicating that these sites are still relatively weak after lamina repair. Patches seemed to grow slowly in size over the course of minutes (Figure 3c). We did not observe PML puncta interacting with these growing patches indicating the most likely source of PMLII is the soluble pool of PMLII. The soluble pool of PMLII is shared between patches and puncta, evidenced by the fact that cells with more patches, have less puncta and vice versa (Figure 3e).

In addition to slow formation of patches, we observed events of sudden collapse and disassembly of PMLII patches. As PMLII patches form at sites of compromised lamina and these sites are known locations of NE rupture, we wondered whether disassembly correlated with NE rupture events. We used GFP-NLS to observe NE rupture as leakage of GFP-NLS into the cytoplasm should be observed upon breaching the NE. Indeed, when observing cells co-expressing mRFP-PMLII and GFP-NLS we observed a decrease of nuclear GFP (white arrow, Figure 4a) followed by the quick collapse of a patch structure into a punctum (arrowhead, Figure 4a) measured as a decrease in patch area (Figure 4b).

**Figure 4:**
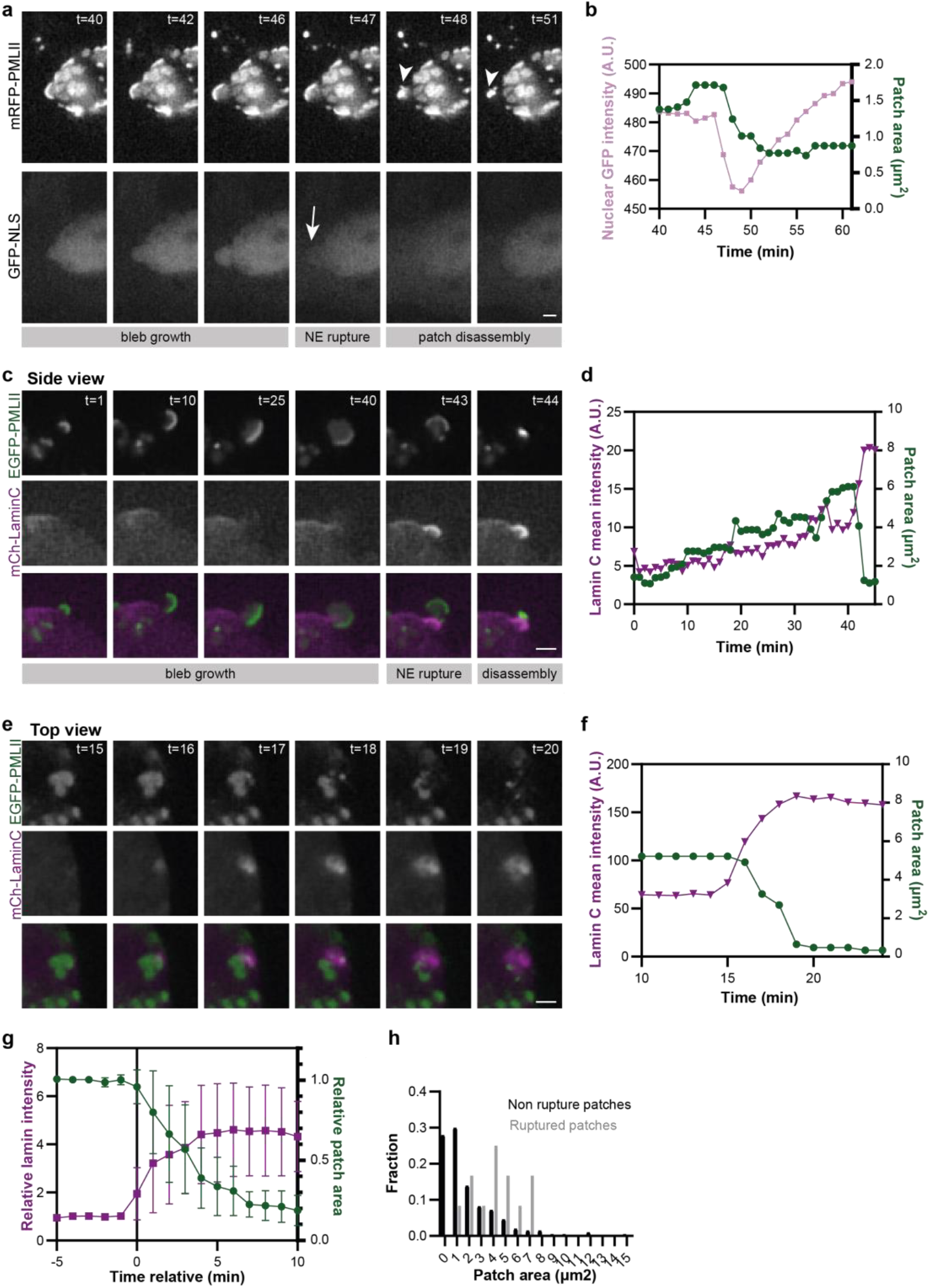
PMLII patches disassemble upon rupture of the NE. **a)** Time-lapse imaging of HeLa cell expressing mRFP-PMLII and GFP-NLS showing a nuclear rupture event indicated by decrease in nuclear GFP signal followed by patch disassembly. Scale bar: 2 μm. **b)** Time-lapse analysis of nuclear GFP intensity and PMLII patch area of and individual NE rupture event showing decrease of nuclear GFP signal preceding patch disassembly. **c)** Example with selected frames from a time-lapse movie of HeLa cells expressing EGFP-PMLII and mCherry-laminC showing bleb growth followed by rupture and initiation of NE repair indicated by lamin C accumulation and subsequent PMLII patch disassembly. Scale bar: 2 μm. **d)** Analysis of lamin C intensity and patch area of example shown in c. **e)** Example with selected frames from a time-lapse movie of HeLa cells expressing EGFP-PMLII and mCherry-laminC representing NE repair associated with PMLII patch disassembly with from a top view. Scale bar: 2 μm. **f)** Analysis of lamin C intensity and patch area of example shown in e. **g)** Average lamin C intensity accumulation over time at the NE associated with patch disassembly. Data was normalized and events were aligned according to the first frame where lamin C signal increase was observed. **h)** Analysis of patch areas from HeLa cells used for timelapse imaging that showed rupture events. N=12 and 193 patches from two independent experiments.

The use of GFP-NLS as a rupture marker often makes it hard to distinguish the exact location of the rupture, especially in smaller events. To more accurately visualize the location of NE rupture events we performed live cell imaging of PMLII together with lamin C. Lamin C has been shown to rapidly accumulate at sites of NE rupture as it is being recruited by the DNA binding protein Barrier-to-autointegration-factor (BAF) to DNA now exposed to the cytoplasmic environment^13,36^. Using lamin C as a rupture marker, we find that NE ruptures occur at sites where PMLII patches are present. We could observe bleb formation events accompanied by PMLII patch growth, after which rapid accumulation of lamin C indicated the occurrence of NE rupture (Figure 4c, d).

### PMLII patches disassemble after initiation of rupture repair

Disassembly of small patches occurs in 1-2 minutes and therefore deciphering the order of events is tricky when observing spontaneous rupture events. Bigger patches however typically take longer to disassemble and provide more detailed spatial information. Disassembly of large patches starts by lamin C accumulation at one side of the patch. The lamin C accumulation spreads while the patch disassembles at these locations (Figure 4e, f). We often observe lamin C accumulation preceding PMLII patch disassembly but never patch disassembly before lamin C accumulation. This indicates that the repair process is initiated before patch disassembly. On average we see full accumulation of lamin C within ∼5 minutes and full patch disassembly within ∼7 minutes (Figure 4g). Finally, we noticed that bigger patches seem more likely to rupture. To confirm a correlation between patch size and the chance of rupturing we analysed patch sizes from cells where we recorded a rupture event. This data indicates that bigger patches are more likely to rupture, although it is not always the biggest patch that ruptures (Figure 4h). Likely other factors contribute such as membrane curvature and local pressure differences.

Whether PMLII is recruited to sites of weakened lamina or whether PMLII plays a role in forming lamin holes is unclear. To look at the consequence of loss of PML and PMLII specifically we turned to knockdown experiments. We observed that loss of all PML, reduced the number of blebs per nucleus (Figure 5a, b). This could suggest that PMLII is important for lamin hole formation or that more general PML bodies play a role in remodelling of the lamina. Indeed, we also found that our PML KO cell line had a lack of lamin holes compared to HeLa cells expressing PML (Figure 5c, d). We then attempted to knockdown PMLII specifically, using previously published PMLII siRNAs^25,37^. However, we found that these siRNAs target PMLII, PMLV and PMLVI, due to use of the PMLII specific Exon 7b in the 3’UTR of these other isoforms. We indeed saw that when targeting PMLII we lost almost all PML. Furthermore, the two siRNAs showed inconsistent effects on NE integrity (Figure 5a, b). Together, we concluded that specific knockdown of PMLII was not achievable.

**Figure 5:**
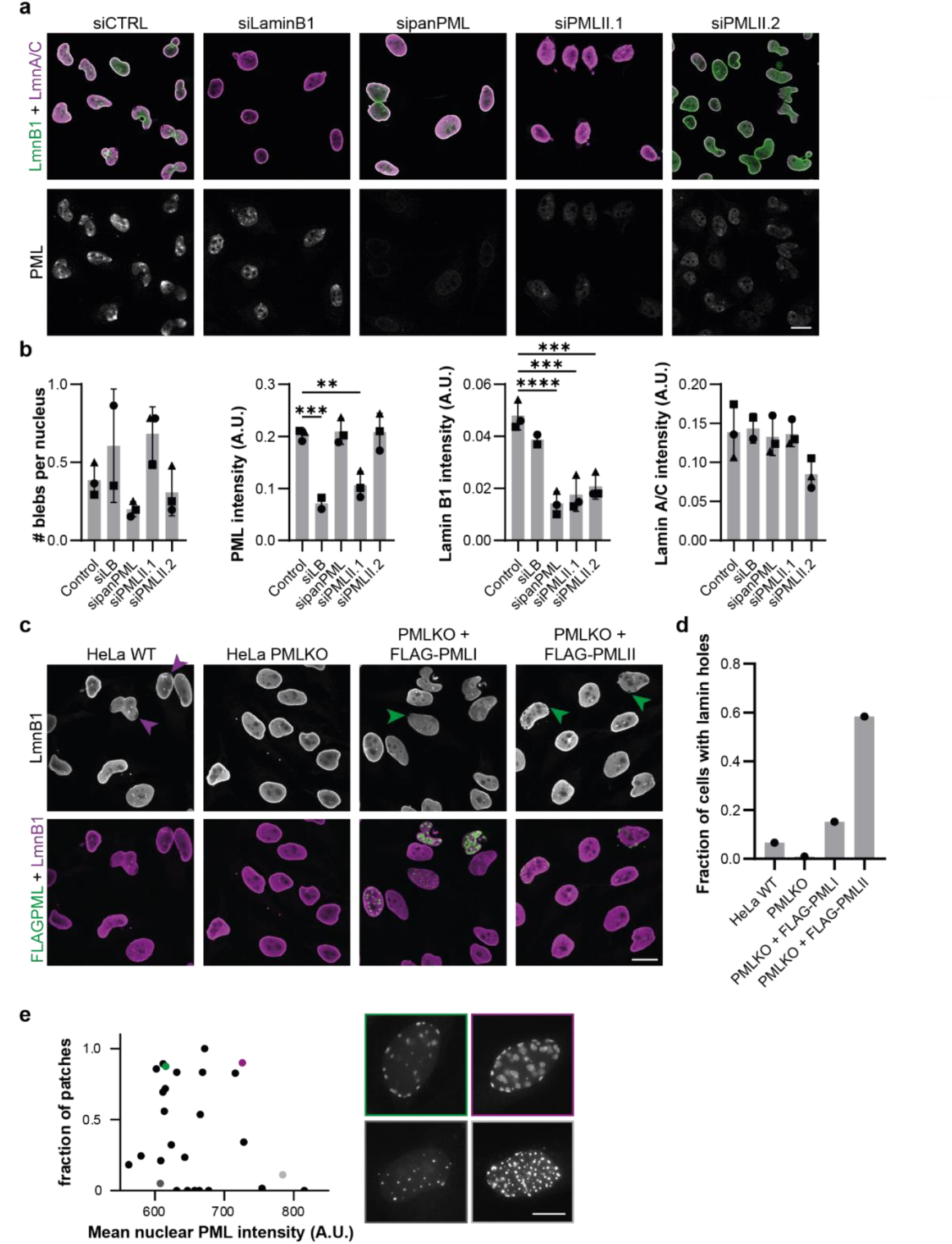
PML is involved in NE maintenance. **a)** Representative, lamin B1 and lamin A/C immunofluorescence images of HeLa cells transfected with lamin B1, PML or two PMLII targeting siRNA. Scale bar: 20 μm. **b)** Quantification of effect of knockdowns in a) on lamin and PML levels and on presence of nuclear blebs. P-value was calculated using a one-way ANOVA test followed by Dunnett’s post hoc test (**P<0.01, ***P<0.001, ****P<0.0001) on n=3 independent experiments (indicated by different symbols), except for siLaminB1 which was n=2. **c)** Representative lamin B1 and FLAG immunofluorescence images of WT HeLa and HeLa PMLKO cells overexpressing FLAG-PMLI and FLAG-PMLII constructs. Purple arrowheads indicate lamin holes in untransfected cells, while green arrowheads indicate holes in cells expressing PML isoforms. Scale bar: 20 μm. **d)** Quantification of effect of presence of PML isoforms on fraction of cells with lamin holes. **e)** Correlation between fraction of patches (#patches/(#patches + #puncta)) and mean nuclear PML intensity in HeLa PMLKO cells overexpressing EGFP-PMLII. n=27 cells. Example cells are indicated by the coloured datapoints and corresponding boxes. Scale bar: 10 μm.

We therefore tried to rescue the lack of lamin holes in PML KO cells by reintroducing PML isoforms. Our data suggests that reintroduction of PMLI can rescue the number of lamin holes in HeLa PML KO cell lines while reintroducing PMLII in a KO background increases the fraction of cells with holes even further, as we observed before (Figure 5c, d). We suggest that PMLII provides structural support when holes in the lamina appear and that PML bodies in general have a role in NE homeostasis. In support of this, we found that there does not seem to be a direct correlation between PMLII nuclear intensity and patch versus puncta formation, suggesting that other events may contribute to the formation of lamin holes (Figure 5e). We thus suggest that PMLII recognizes lamin defects, that are caused by other cellular mechanisms.

## Discussion

NE rupture is a major remodelling event with possible detrimental effects on cellular and organismal health. Quick repair reducing exchange of material between the nuclear interior and exterior is therefore important to minimize such consequences. In this study, we observed that PMLII can form patch like condensates at NE sites with weakened lamina. Emerging evidence indicates important roles for interactions between biomolecular condensates and membrane-bound organelles in cellular organization^19–22^. Our observations lead to a model where PMLII patch assembly occurs at sites of weakened lamina (Figure 6). At some point, these local weak spots in the NE can rupture and the repair machinery, including lamin C, is quickly recruited. We observed that the start of rupture and repair precedes the collapse of the PMLII patch. We hypothesise that the PMLII patch provides structural integrity to the site of rupture preventing more extensive ruptures of the membrane and providing stability for repair to occur. This suggested that the role for PMLII condensates in stabilizing the rupture site is similar to a recently reported role for stress granules in stabilization of damaged endolysosomes^22^ and could point towards a more general principle of stabilization of cellular membranes by condensates. While stress granules seem to rapidly assemble on ruptured membranes, we observe a slow accumulation of PML condensates at weakened nuclear envelope sites before membrane rupture is observed. This suggests that PML condensates might also play a role in preventing rupture to occur although we never observed PMLII patch disassembly without a rupture event. Whether this means that lamin holes cannot repair without rupture events or whether we simply do not observe these events in our conditions is unclear. We are observing micron sized lamin holes, in cancer cells under conditions of overexpression. We speculate that under more physiological conditions the repair of small lamin holes would be possible as it would be beneficial for the cell to be able to repair small lamin defects without the need of a rupture event. Interestingly, PMLII has been linked to nuclear lipid synthesis and lipid droplets^25,38^. Whether local lipid synthesis plays a role in repair of NE membrane ruptures or more general NE integrity maintenance and remodelling remains to be investigated.

**Figure 6:**
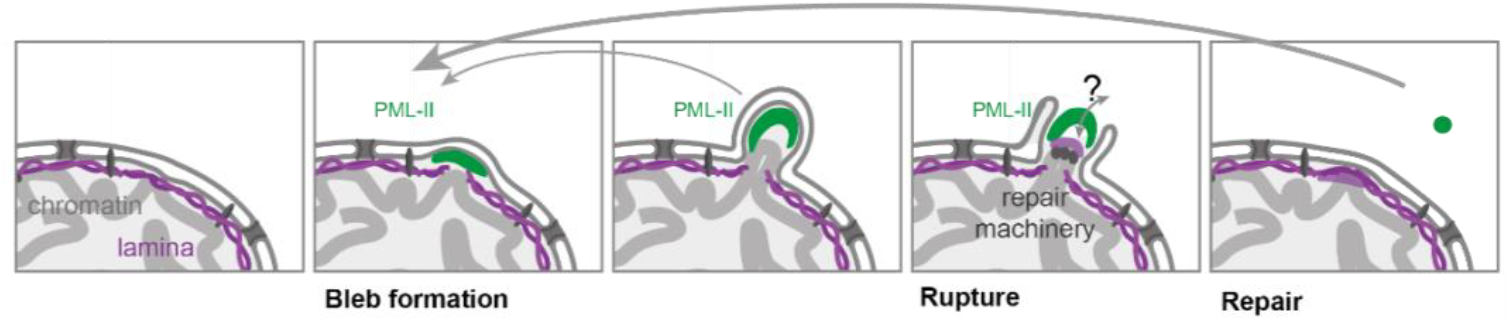
Schematic representation of PMLII function in a NE rupture event providing support for membrane repair to be initiated.

We noticed that overexpression of PMLII leads to formation of a large number of patches and lamin holes, not typically observed when looking at endogenous PML. We showed that the resulting high levels of PMLII compared to other isoforms likely plays a role as more PMLII is now available for patch formation instead of recruitment to bodies. This also poses the question whether PMLII might play a role in formation of lamin holes. Our data indicates a general role for PML bodies in lamina maintenance as loss of all PML leads to a decrease in lamin holes and reintroduction of PMLI can rescue this. Sumoylation has been found to play roles in regulation of nuclear lamins^39^ and NE dynamics more generally^40–44^. For example, in yeast it was found that temporally controlled INM sumyolation plays a role in coordinating NE biogenesis during mitosis^45^. We therefore speculate that loss of PML bodies, that constitute a major nuclear hub for sumoylated proteins, leads to deregulation of NE sumoylation events and therefore affect NE maintenance. We suggest that a delicate balance between PMLII at bodies versus interacting with the NE ensures that PMLII in normal situations does not induce massive holes in the lamina, but we do not rule out a role for PMLII in lamina maintenance for example in events where local lamina remodelling is required.

We showed that the amphiphatic helices in PMLII are important for localization to NE patches. Amphipathic helices are often found to interact with membranes by partial insertion into one leaflet of the membrane. This insertion disturbs local headgroup packing and therefore can lead to curvature sensing or inducing properties. However, we would expect an opposing curvature on the membrane than we observe. Scaffolding is another mechanism by which membrane curvature can be sensed or induced^33^. Specifically, attractive forces between intrinsically disordered domains have been shown to drive concave bending^46^. We therefore envision that PMLII condensation combined with membrane affinity provided by the α-helices senses/induces membrane curvature leading to localization of PMLII to NE blebs. Interestingly, curvature induced lamin dilution has been a proposed mechanism for loss of lamin at NE blebs^47^. Whether PMLII condensates recognize a specific lipid composition at these curved sites, whether PMLII is recruited by other partners or whether merely the lack of a dense lamin network is what allows PMLII to localize to lamin holes remains unclear.

## Materials and Methods

### Cell culture

HeLa, U2OS and hTERT fibroblasts were cultured in Dulbecco’s modified Eagle’s medium containing 10% foetal calf serum and penicillin/streptomycin and were maintained at 37°C and 5% CO_2_. hTERT fibroblast cell line was derived from AG10803 immortalized with SV40LT and TERT (gift from Carlos López-Otín).

### Generation of PML knockout cells

We used an All-in-One Cas9^D10A^ nickase vector with enrichment by fluorescence-activated cell sorting to enrich for transfected cells^48^. A pair of guide RNAs (sgRNAs) was designed to target the DNA on opposing strands on Exon2 of the PML gene. The sgRNAs were then cloned into the All-in-One Cas9^D10A^ nickase vector using DNA oligos (Sigma-Aldrich) using the BsaI and BbsI recognition sites. The sgRNA sequences: TGTCTGCACACGCTGTGCTC and TGGCTTCCGCCTGGCATTGC were used for CRISPR targeting. Cells were transfected with the All-in-One vector using Transit2020. GFP positive cells were sorted as a polyclonal population for manual seeding into 96-well plates. After expansion, cells were seeded on coverslips and immunofluorescence was performed to screen for successful knockouts.

Immunoblotting was performed to confirm loss of PML protein. HeLa WT and HeLa PML KO clone57 cells were lysed in Laemmli buffer (4% SDS, 20% glycerol, and 120 mM Tris-HCl [pH 6.8]) and then incubated for 5 min at 95°C. The DNA was sheared by syringing the lysates 10 times through a 25-gauge needle. Absorbance at 280 nm was measured (NanoDrop; Thermo Fisher Scientific) to determine protein concentration. Samples were prepared using Protein Sample Loading Buffer (LI-COR, #928-40004) and DTT (final concentration 25 mM). Lysates were collected on different passages (n=3). 40 ug of protein was loaded for all samples. Proteins were separated using NuPAGE 4-12% Bis-Tris gels (ThermoFisher) and NuPAGE MOPS SDS running buffer (Thermo Fisher, #NP0001) and transferred to nitrocellulose membranes for immunoblotting. Membranes were blocked in 5% milk PBS and incubated overnight at 4°C with primary antibodies. Next day, membranes were incubated for 1 h at room temperature with IRDye-conjugated secondary antibodies (LI-COR) and scanned on an Odyssey imaging system. The following primary antibodies were used: mouse anti-Tubulin (Sigma Aldrich, #T9026, 1/1000), rabbit anti-PML (Abcam, #ab179466, 1/1000).

### Plasmid transfection and constructs

Transfections of plasmid constructs were performed using Trans-IT 2020 reagent (#MIR 5400; Mirus Bio) following the manufacturer’s protocols. 24 h after transfection, experiments were performed. mRFP1-PMLI and PMLII containing plasmids were a kind gift from Peter Hemmerich and were subcloned into a pEGFP-C1 plasmid (Clontech). FLAG-PMLI and FLAG-PMLII plasmids were subsequently generated using Gibson Assembly. PMLII mutants in pEGFP-C1 vector were also generated using Gibson Assembly by assembly of a SalI/XhoI digested pEGFP-C1 with the desired mutant fragments using pcr amplification.

PMLII full length is 829 amino acids and the predicted helices spanned residue 649-674 (helix1) and 740-762 (helix2). We therefore generated the following mutants: PMLIIΔ1: 1-764; PMLIIΔ2: 1-738; PMLIIΔ3: 1-676; PMLIIΔ4: 1-648; PMLIIΔhelix1: deletion of residues 649-674; PMLIIΔhelix2: deletion of residues 740-762; PMLIIHelix1* (charge swap): V651T, I655E, L659D; PMLIIHelix2* (charge swap): L743D, L747D, F651D, M757E; sumoylation mutant: PMLIIK160R; B1Box muant: PMLIIF158E. GFP-NLS was described previously^49^ and contains cycle3GFP fused to EGFP, an NLS, and a second EGFP in a pcDNA backbone (kind gift from Emily Hatch).

The pICE-mCherry-laminC was constructed by cloning in the HindIII and XhoI digested pICE (deposited on Addgene, plasmid #46960, PMID:23897892), the mCherry sequence, amplified by PCR from pmCherry-C1 (Clontech) using mCh-F GCCAAGCTTACCGGTCGCCACCATGGTGAGC and mCh-R CCGAGATCTGAGTCCGGACTTGTACAGC primers and digested by HindIII and BglII, and the lamin C human sequence, amplified by PCR from the lamin A cDNA (IMAGE clone #4863480) using LmA-F GCGAGATCTATGGAGACCCCGTCCCAGCG and LmC-R GGCCTCGAGTTAGCTGCTGCAGTGGGAGCCGTGG primers and digested by BglII and XhoI.

### siRNA knockdown

The following siRNA oligonucleotides were purchased from Sigma-Aldrich: sipanPML: 5′-AGAUGCAGCUGUAUCCAAGdTdT-3′, siPMLII.1: 5′-CAUCCUGCCCAGCUGCAAAdTdT-3′, siPMLII.2: 5’-GUGGCUCAACAACUUUUUUdTdT-3’. For knockdown of lamin B1 Dharmacon siGENOME SMART pool targeting human LMNB1 (Cat #M-005270-01-0005) was used. MISSION siRNA Universal Negative Control #1 (#SIC001; Sigma-Aldrich) was used as the control siRNA. 30 pmol of siRNA were transfected into cells in 12-well plates using Lipofectamine RNAiMAX (#13778075; Thermo Fisher Scientific) following the manufacturer’s instructions. After 72h cells were fixed for subsequent immunofluorescence.

Lamin A/C, lamin B1 and PML levels in knockdown cells were measured by automated detection of nuclei using Cellprofiler object detection in the DAPI channel. Blebs were detected as previously described^50^. In brief: Chromatin object was identified based on DAPI staining. A lamin B1 object was identified, and subtraction of both areas identifies the bleb object as it is devoid of lamin B1. Subsequently, bleb areas are filtered to exclude small, identified regions or pixels at the edge of the nuclei. Size, shape and number of bleb areas are measured.

### Immunofluorescence

Cells were fixed with 4% PFA for 10 minutes at room temperature. Cells were washed in PBS, permeabilized using 0.2% Triton-X100 for 15 minutes, and blocked using 2% bovine serum albumin (BSA, Sigma-Aldrich, #A7638) in PBS for 30 min. Cells were incubated overnight at 4°C or for 1 h at RT in 2% BSA PBS containing primary antibody. Cells were washed using PBS and incubated for 1 h at room temperature with secondary antibody in 2% BSA PBS. Cells were washed in PBS and mounted using Prolong Diamond (Thermo Fischer, #P36965). The following primary antibodies were used: mouse anti-lamin A/C (Santa Cruz, sc-376248, 1/500), mouse anti-lamin B1 (Santa Cruz, sc-365214, 1/500), mouse anti-FLAG (Sigma Aldrich, F1804, 1/1000), rabbit anti-PML (Abcam, ab179466, 1/500) and rabbit anti-sp100 (Atlas antibodies, HPA016707, 1/500). Confocal images were taken on a Leica Stellaris 5 confocal laser scanning microscope using a 63x oil objective (PL APO CS2, 1.40 NA). Widefield images were taken on a Nikon AX R Confocal microscope in widefield mode using a 60x oil immersion objective (PLAN APO, 1.4 NA) and a Nikon DS-Qi2 Camera.

### Image processing and analysis

Images were analysed using ImageJ or Cellprofiler V8 as indicated. Data were transferred to Microsoft Excel for further calculations and subsequently plotted and analysed using GraphPad Prism 10.

#### PMLI/II ratios

Nuclei were automatically detected using Cellprofiler by object detection using DAPI staining. Nuclear PMLI and PMLII mean intensities were then measured and the ratio was determined. Condensate types were then manually assigned to the detected nuclei based on visual inspection of PMLII localization. Cells on the edge of the image, that showed very severe abnormalities or had very high levels of PML expression resulting in aggregation were discarded from the analysis.

#### Lamin B1 and sp100 levels at bodies and patches

Custom Cellprofiler pipelines were used for automated detection and analysis of patches and bodies. In short: PML condensates were detected based on the EGFP channel and were discarded when outside of the nucleus based on nuclei detection in the DAPI channel. Size and shape of detected objects were measured and used to discriminate between patches and bodies. Finally, lamin B1 and sp100 intensities of patches and bodies were measured using the filtered objects. Background substraction was performed based on intensity measurements outside the cell. For relative lamin B1 levels, the levels of lamin B1 mean intensity at patches was divided by the mean intensity of the entire nucleus. Per nucleus means were calculated which were then used to determine experimental means for 3 biological repeats.

#### Condensate types

Condensate types were manually assigned. Cells on the edge of the image, that showed very severe nuclear abnormalities or had very high levels of PML expression resulting in aggregation were discarded from the analysis.

#### Patches versus puncta numbers

HeLa PMLKO cells expressing EGFP-PMLII were imaged on a widefield Nikon Ti2 and analysed manually by counting the number of puncta and patches. Additionally, mean nuclear EGFP fluorescence was measured by drawing an area encompassing the nucleus using lamin B1 staining.

### Live cell imaging and analysis

HeLa cells were seeded in 4 well μ-slides (#80426, Ibidi). Timelapse imaging was performed 24hr after transient transfection on a Nikon AX R Confocal microscope in widefield mode using a 40x water immersion objective (APO, 1.15 NA) and a Nikon DS-Qi2 Camera.

#### Analysis GFP-NLS, lamin C and patch shrinkage

Patch size was measured by manual drawing of areas and intensities were measured in this area. When patch started shrinking and falling apart the biggest area describing the patch was measured. In addition, a larger oval surrounding the patch was used to measure lamin C recruitment to the patch location even after patch disassembly. Background subtraction was performed by measuring an area outside the nucleus. GFP-NLS levels in the nucleus were measured by drawing a circular ROI in the nucleus and measuring mean GFP intensity over time.

#### Patch area and rupture chance

Live cell imaging data of HeLa cells expressing EGFP-PMLII and mCherry-laminC showing rupture events were analysed. The frame before an observed rupture event was used to analyze patch area. Using LabKit Plugin in ImageJ, automated segmentation of PMLII condensates was performed. Area of the detected regions was subsequently measured, and these were manually filtered based on condensate type only leaving patch structures. Detected patch sizes were either classified as being the site of a future rupture event or not.

#### Curvature

Curvature of PMLII patches and the nucleus were analysed based on live cell imaging data. Patches that clearly were in the focal plane on the side of the nucleus were analysed by drawing a line and analysing curvature in 2D using the Kappa-curvature analysis plugin in ImageJ. Nuclear curvature was estimated based on mCherry-laminC imaging.

### Fluorescence recovery after photobleaching

For FRAP analysis of condensate dynamics, HeLa FlpIn EGFP-PMLII cells were used. To generate stable cell lines, HeLa FlpIn TetR cells were transfected with pOG44 and pcDNA5/FRT/TO-EGFP-PMLII using Lipofectamine 3000 (Thermo Fisher, L3000001). Selection was performed using 200 ug/mL Hygromycin. After selection cells were maintained in 100 ug/mL Hygromycin. EGFP-PMLII expression was induced by 100 ng/ul Dox addition for 24-30hr before the start of the experiment. Condensates were imaged on a Nikon SoRa spinning disk confocal in normal confocal mode using a 100x oil objective (CFI SR HP Apo TIRF, NA 1.49) and an ORCA-fusion BT digital camera (Hamamatsu, C15440). Cells were imaged at 37°C, 5%CO2 using a Tokai Hit STX Stage Top incubator. An opti-microscan FRAP scanner was used to bleach condensates using a Oxxius LBX-488 laser. Cells were imaged for 7 min every 5s to monitor recovery. Intensity values were measured using ImageJ and normalized with pre-bleaching intensities after background correction. Frames were puncta clearly moved out of focus were discarded.

### Structure prediction and analysis

The AlphaFold2 model of the PMLII C-terminus was predicted using default parameters in ColabFold [ref PMID: 35637307]. Images were generated using PyMOL (Schrödinger, LLC). The helical wheel representation of the amphipathic sequence was generated using the HeliQuest server [ref PMID: 18662927].

## Acknowledgements

D Larrieu was funded by a Sir Henry Dale Fellowship jointly funded by the Wellcome Trust and the Royal Society (Grant Number 206242/Z/17/Z). AFJ Janssen was supported by a Leverhulme Trust Early Career Fellowship and the Isaac Newton Trust. O Knowles was funded by the University of Cambridge Pharmacology Department. E Spruijt acknowledges financial support from a Vidi grant from the Netherlands Organization for Scientific Research (NWO, Vidi. 193.089). JE Deane is supported by a Wellcome Trust Senior Research Fellowship (219447/Z/19/Z). We would like to thank Dr. Janet Kumita for fruitful discussions on the project. We would also like to acknowledge Cambridge Advanced Imaging Centre, the Gurdon Institute Imaging Facility and Radboud University Light microscopy core-facility for instrumentation and advice.

## Author contributions

A.F.J.J. and D.L. designed research. A.F.J.J. and O.K. created reagents. S.B. constructed the mCherry-laminC plasmid. J.E.D. performed structure prediction and analysis. A.F.J.J. performed experiments, analysed data and prepared figures. A.F.J.J. wrote the original draft. A.F.J.J., O.K., S.B., J.E.D., E.S. and D.L. reviewed and edited the paper. A.F.J., E.S. and D.L. acquired funding. D.L. and E.S. supervised the project. All authors have read and agreed to the final version of the manuscript.

