## Supplemental Figures for "Patching up the nucleus: a novel role for PMLII in nuclear envelope rupture repair"

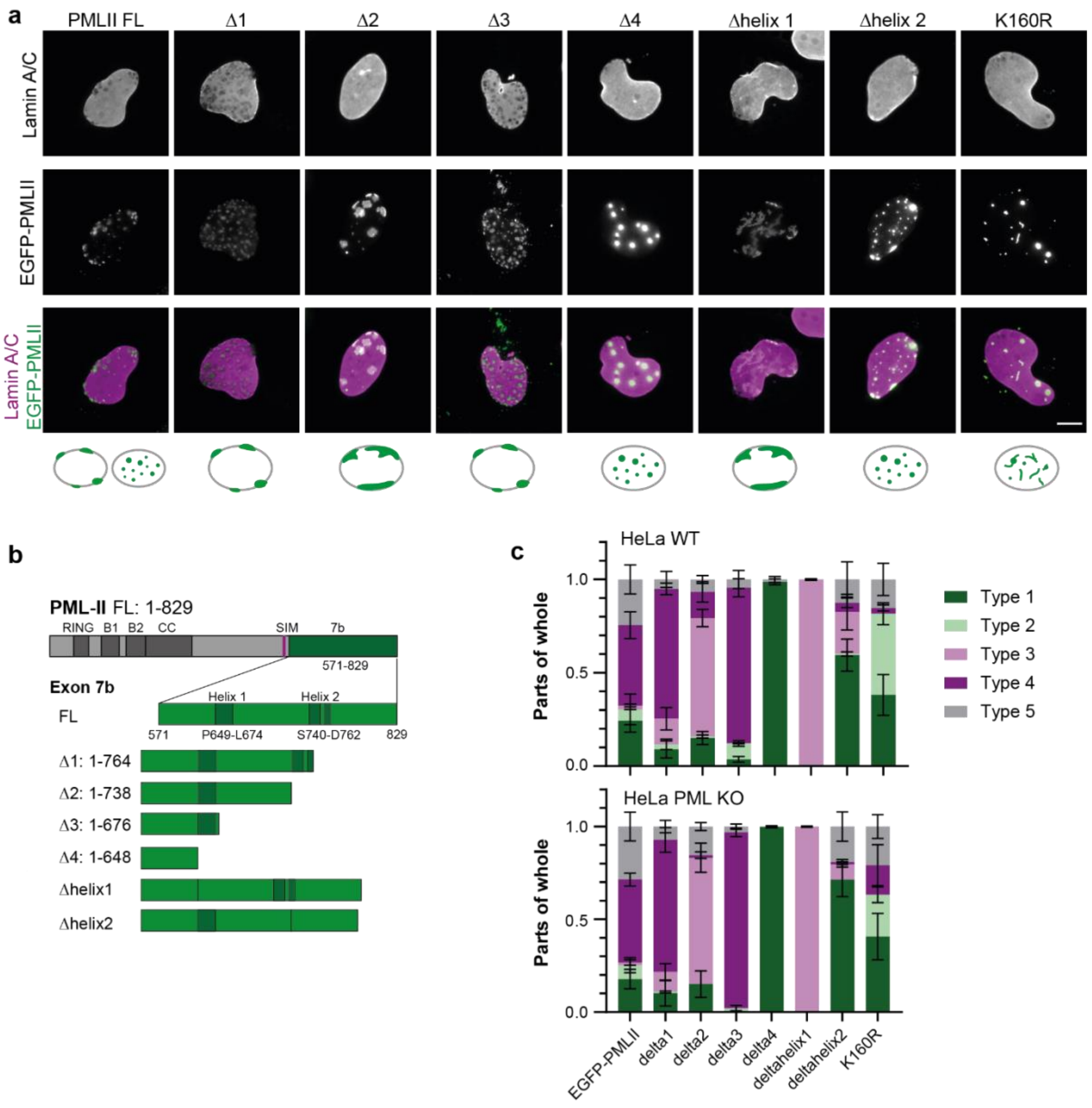


**Figure S1: Effect of PMLII C-terminal truncations and mutations on condensate formation. a)** Lamin B1 immunofluorescence images of HeLa cells overexpressing EGFP-PMLII WT and mutants. Examples show the most abundant condensate type according to manual classification. Scale bar: 10 µm. **b)** Schematic representation of PMLII organization with the common N-terminal RBCC domain and unique C-terminus (exon 7b) and the generated mutants. **c)** Quantification of fractions of HeLa WT and PML KO cells overexpressing EGFP-PMLII WT and indicated mutants showing different types of condensates. Mean +/- sd indicated from N=3 independent experiments.

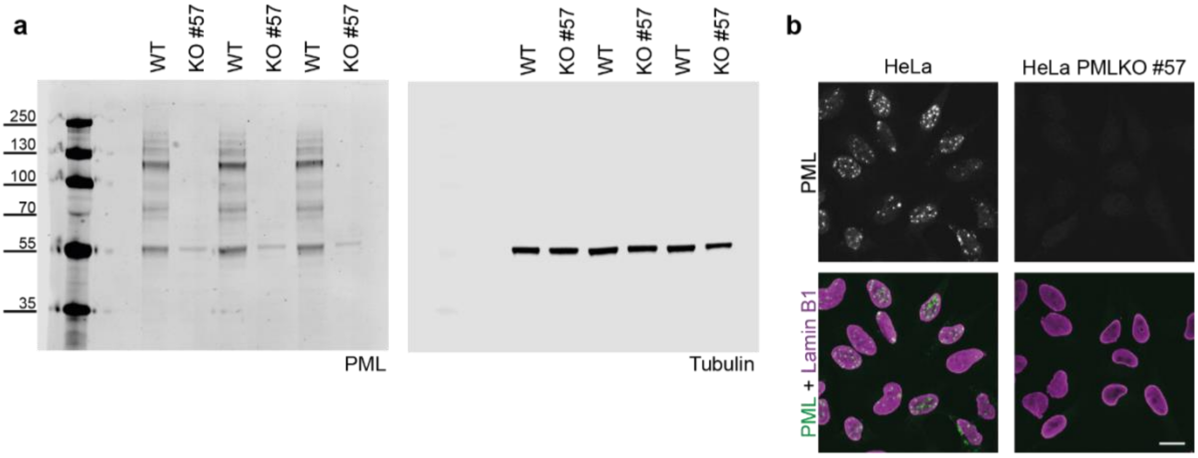


**Figures S2: PML knockout HeLa cells. a)** PML and Tubulin immunoblot of whole cell lysate from HeLa WT and PMLKO cells. 3 samples were collected at different timepoints. **b)** PML and lamin B1 Immunofluorescence images of HeLa WT and PMLKO cells. Scale bar, 20 μm.
